# Cholesterol binding to VCAM-1 promotes vascular inflammation

**DOI:** 10.1101/2024.09.17.613543

**Authors:** John P. Kennelly, Xu Xiao, Yajing Gao, Sumin Kim, Soon-Gook Hong, Miranda Villanueva, Alessandra Ferrari, Lauri Vanharanta, Alexander Nguyen, Rohith T. Nagari, Nikolas R. Burton, Marcus J. Tol, Andrew P. Becker, Min Jae Lee, Elina Ikonen, Keriann M. Backus, Julia J. Mack, Peter Tontonoz

## Abstract

Hypercholesterolemia has long been implicated in endothelial cell (EC) dysfunction, but the mechanisms by which excess cholesterol causes vascular pathology are incompletely understood. Here we used a cholesterol-mimetic probe to map cholesterol-protein interactions in primary human ECs and discovered that cholesterol binds to and stabilizes the adhesion molecule VCAM-1. We show that accessible plasma membrane (PM) cholesterol in ECs is acutely responsive to inflammatory stimuli and that the nonvesicular cholesterol transporter Aster-A regulates VCAM-1 stability in activated ECs by controlling the size of this pool. Deletion of Aster-A in ECs increases VCAM-1 protein, promotes immune cell recruitment to vessels, and impairs pulmonary immune homeostasis. Conversely, depleting cholesterol from the endothelium *in vivo* dampens VCAM-1 induction in response to inflammatory stimuli. These findings identify cholesterol binding to VCAM-1 as a key step during EC activation and provide a biochemical explanation for the ability of excess membrane cholesterol to promote immune cell recruitment to the endothelium.

## Introduction

Cytokines, pathogens, and other pro-inflammatory agents ‘activate’ endothelial cells (ECs), conferring on them enhanced ability to attract and bind leukocytes^1^. Leukocyte recruitment to ECs is a critical step in the propagation and resolution of inflammation, wound healing, and thrombosis^2^. Failure to properly control EC-leukocyte interactions is linked to the etiology of diseases including atherosclerosis, reperfusion injury, inflammatory bowel disease, and acute lung injury^2^. Leukocyte binding to ECs is facilitated by plasma membrane (PM)-embedded adhesion molecules, including vascular cell-adhesion molecule 1 (VCAM-1)^3,4^. The mechanisms by which PM lipid composition influence EC adhesiveness are poorly understood.

Most unesterified cellular cholesterol is concentrated in the PM^5^. Cholesterol in the PM exists in at least two forms: a pool that is sequestered by phospholipids (primarily sphingomyelin; SM) and a more mobile ‘accessible’ cholesterol pool^6^. PM cholesterol becomes ‘accessible’ for interactions with transporters or other proteins when it is present in amounts that exceed the capacity of local membrane phospholipids to sequester it^6^. Accessible cholesterol has more chemical potential than cholesterol complexed with phospholipids due to its greater ability to enter different metabolic pathways or modulate protein function^7^. For example, the PM accessible cholesterol pool influences the rate of cellular cholesterol biosynthesis and uptake because its transfer to ER membranes inhibits SREBP-2 processing^8^. Accessible cholesterol transport to the ER also enables the production of cholesteryl esters, oxysterols, bile acids, and steroid hormones.

The nonvesicular cholesterol transport proteins Aster-A, -B, -C (encoded by *Gramd1a*, *Gramd1b* and *Gramd1c,* respectively) mediate accessible cholesterol movement from the PM to the ER in mammalian cells^9,10^. Asters are anchored to the ER by a single-pass transmembrane domain (TMD), and they form contacts with cholesterol-enriched PMs via an N-terminal GRAM domain^9^. Asters are important for PM-ER cholesterol transport in tissues that store or secrete large amounts of cholesteryl esters, including the adrenal, liver, intestine, and ovary^9,11–13^. However, whether the ability of lipid trafficking pathways to enrich or deplete organelle membranes of specific lipids can modulate membrane protein function in other physiological settings remains to be explored.

Hypercholesterolemia promotes leukocyte binding to the endothelium^14–16^, and is associated with the development of inflammatory disorders including atherosclerosis, psoriasis, and psoriatic arthritis^17,18^. Cytokines released from cholesterol-laden foam cells within the artery wall are known to promote the transcription of adhesion molecules in the setting of hypercholesterolemia^19,20^. However, several lines of evidence suggest that excess cholesterol also acts directly on the endothelium to increase its susceptibility to activation and dysfunction. For example, the removal of low-density lipoprotein (LDL) cholesterol from the blood of people with hypercholesterolemia by apheresis acutely improves endothelium-dependent vasodilation^21^ and lowers markers of EC activation^22^. Despite these links, mechanistic insight into how cholesterol accumulation increases EC adhesiveness is lacking. Our understanding of the repertoire of PM proteins whose abundance or activity is modulated by interaction with cholesterol is also incomplete.

In the current study we utilized cholesterol-mimetic photoaffinity probes in combination with mass spectrometry-based proteomics to map cholesterol-protein interactions in primary human ECs. We find that cholesterol binds directly to VCAM-1, thereby preventing its ubiquitination and proteasomal degradation. During EC activation and in the setting of hypercholesterolemia, VCAM-1 is bound and stabilized by PM accessible cholesterol. Expanding the accessible cholesterol pool in the PM of ECs *in vivo* by genetic deletion of Aster-A increases VCAM-1 on the endothelium, promotes immune cell adhesion to vessels, and causes pulmonary inflammation. Conversely, Aster-A overexpression or cholesterol extraction from the EC membrane blunts VCAM-1 induction in response to inflammatory stimuli. These findings identify accessible membrane cholesterol as a physiological and pathophysiological modifier of vascular inflammation.

## Results

### Cholesterol directly binds to VCAM-1

To identify cholesterol-interacting proteins in human ECs, we synthesized a cholesterol-mimetic photoaffinity probe named NBII-165 (**Fig. 1a**). The probe consists of an intact cholesterol backbone with a photoreactive diazirine group on the alkyl tail for ultraviolet (UV) light–induced cross-linking to interacting proteins and a terminal alkyne group for enrichment by copper(I)-catalyzed azide–alkyne cycloaddition (CuAAC) or “click chemistry” (**Fig. 1a**). NBII-165 suppressed SREBP-2 pathway targets in human umbilical vein ECs (HUVECs) to a similar extent as cholesterol or the previously characterized cholesterol mimetic probe KK-174^23^ (**Fig. 1b and Extended Data Fig. 1a**), confirming that NBII-165 mimics cholesterol in a well-validated biological assay.

**Fig. 1.**
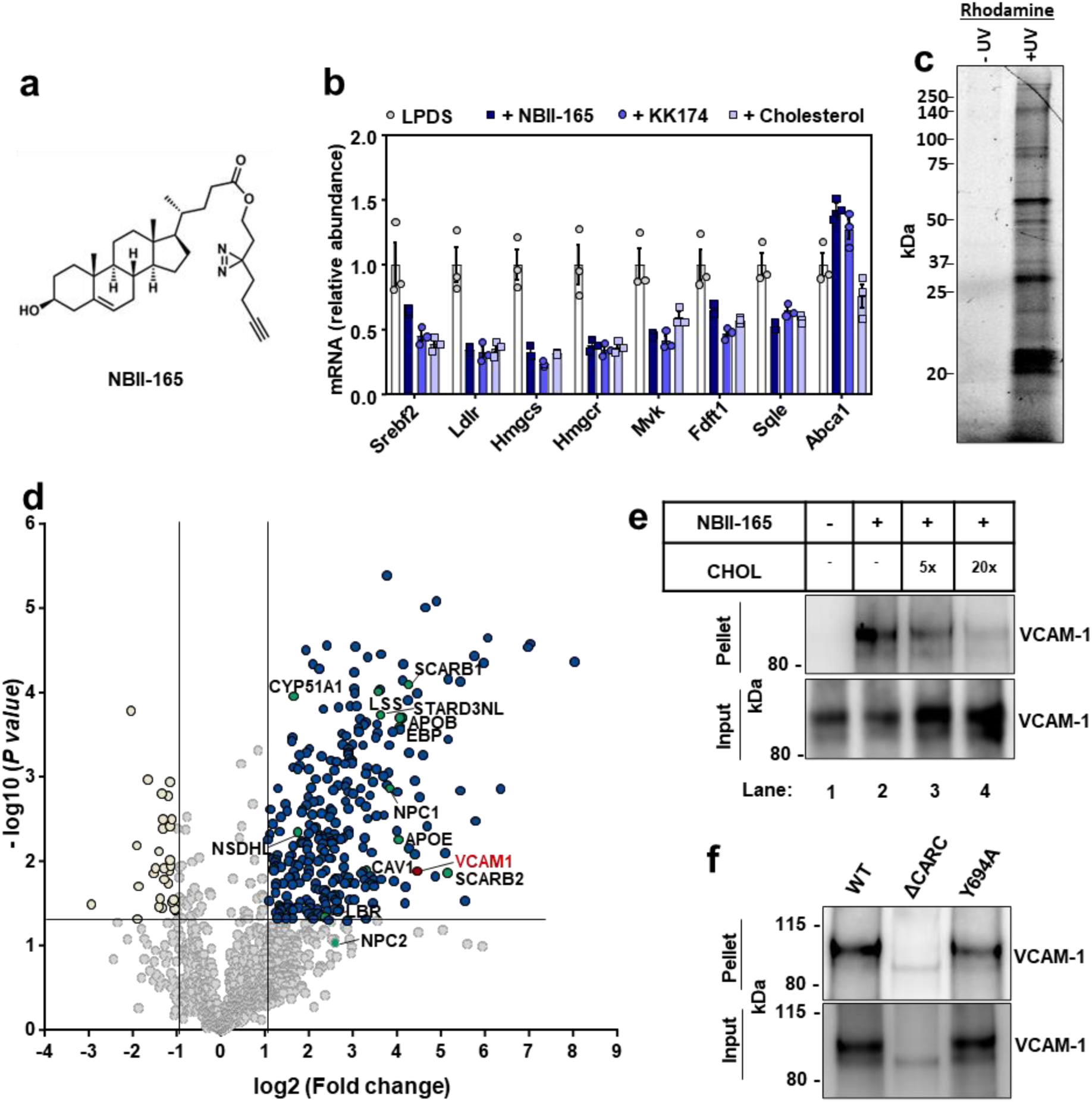
Cholesterol-mimetic probes identify VCAM-1 as a cholesterol binding protein in primary human ECs. (a) Structure of NBII-165 probe. (b) HUVECs were depleted of cholesterol overnight in LPDS containing simvastatin. Cells were then loaded with 35 µM NBII-165, KK-174 or cholesterol complexed to methyl-beta cyclodextrin for 4 h before collection and assessment of SREBP-2 targets by qPCR. (c) Rhodamine-azide signal in HUVEC lysates. HUVECs were incubated with 10 μM NBII-165 probe for 1 h, with and without 365 nm UV irradiation before attachment of a rhodamine-azide fluorophore by click chemistry and separation of proteins by sodium dodecyl sulfate (SDS)-polyacrylamide gel electrophoresis (PAGE). The in-gel rhodamine signal was visualized with a fluorescent imager. (d) Volcano plot showing proteins that were detected by mass spec after immunoprecipitation of NBII-165-bound proteins. Dark blue dots indicate significantly enriched proteins. Yellow dots indicate proteins that were significantly lower in the UV exposed samples. Green dots indicate known sterol binding proteins. Burgundy dot indicates VCAM-1. (e) Competition assay showing that cholesterol competes with NBII-165 for binding to VCAM-1 in HUVECs stably overexpressing human VCAM-1. Input shows VCAM-1 detected in whole cell lysates prior to immunoprecipitation and pellet shows VCAM-1 detected after streptavidin immunoprecipitation of probe bound proteins. (f) Immunoprecipitation of WT VCAM-1 or mutant versions of VCAM-1 either lacking the CARC motif or with a tyrosine for alanine mutation at amino acid 694 after incubating HUVECs with KK-174 followed by UV crosslinking. Data are represented as mean ± SEM.

For probe interaction experiments, we delivered NBII-165 to HUVECs for 1 h before UV crosslinking to interacting proteins. The UV-dependence of NBII-165 labeling was confirmed by click-conjugation of a rhodamine-azide tag to probe-bound proteins before in-gel imaging of the fluorescent rhodamine signal (**Fig. 1c**). Competition assays showed that protein labeling events in cells treated with either NBII-165 or KK-174 could be dose-dependently reduced by co-incubation with excess cholesterol (**Extended Data Fig. 1b**), with no change to total protein loading (**Extended Data Fig. 1c**). To facilitate identification of cholesterol-interacting proteins in activated ECs, we treated cells with lipopolysaccharide (LPS) 6 h prior to NBII-165 delivery. A negative control dataset was generated by administering the probe to LPS-activated ECs that did not receive subsequent UV light exposure. After cell collection and lysis, NBII-165-labeled proteins were conjugated to an azide-biotin tag by click chemistry^24^ before streptavidin affinity enrichment and on-bead trypsin digestion.

Mass spectrometry identified 289 proteins that were enriched (log_2_ fold change > 1 and *P* value < 0.05) in the UV-exposed samples compared to control samples (**Fig. 1d**). The list included multiple known sterol-binding proteins, including those involved in cholesterol uptake (SCARB1), cholesterol transport (NPC1, APOE, APOB, STARD3NL, SCARB2), and cholesterol synthesis (LSS, EBP, NSDHL, CYP51A1, LBR). The screen also identified Caveolin-1 (CAV1), which has long been recognized to bind cholesterol in ECs^25^. Unexpectedly, the leukocyte adhesion molecule VCAM-1 emerged as a top candidate cholesterol-binding protein (**Fig. 1d**). Streptavidin pull-down experiments after NBII-165- or KK-174-labeled proteins were conjugated to an azide-biotin tag by click chemistry confirmed that VCAM-1, as measured by western blot, was enriched from ECs after UV irradiation in a probe-dependent manner (**Fig. 1e and Extended Data Fig. 1d**). Interaction between NBII-165 or KK-174 and VCAM-1 was reduced in the presence of excess cholesterol (**Fig. 1e and Extended Data Fig. 1d**). While incubating cells with excess cholesterol reduced the amount of VCAM-1 that was pulled-down with the probes as expected due to competition, we noted that the total amount of VCAM-1 in the whole-cell input was increased by cholesterol delivery (**Fig. 1e and Extended Data Fig. 1d**). This observation suggested that cholesterol binding might stabilize VCAM-1.

VCAM-1 consists of a large extracellular domain that binds to leukocytes, a single-pass TMD that spans the PM, and a short carboxy-terminus cytoplasmic tail. Interestingly, the amino acid sequence of human and mouse VCAM-1 TMD contains a predicted CARC motif (K/R–X_1–5_–Y/F–X_1–5_–L/V; **Extended Data Fig. 1e**). Such sequences are present in other cholesterol-binding proteins^26^. Full-length human VCAM-1 was efficiently pulled-down with KK-174 after UV crosslinking and biotin-azide conjugation by click chemistry (**Fig. 1f**). However, a mutant VCAM-1 with a tyrosine for alanine substitution in the middle of the CARC motif (Y694A) was retrieved less efficiently than the wild-type (WT) protein (**Fig. 1f**). A mutant VCAM-1 protein lacking all thirteen amino acids comprising the CARC motif, although poorly expressed, showed minimal interaction with KK-174 (**Fig. 1f**). These data suggest that cholesterol interacts with the CARC motif in the TMD of VCAM-1.

### Cholesterol binding stabilizes VCAM-1

To further examine the relationship between cholesterol and VCAM-1 protein levels on ECs, we manipulated cholesterol availability in cultured cells. ECs grown in cholesterol-depleted media (lipoprotein-deficient serum (LPDS) and simvastatin) showed lower cell surface binding of the accessible PM cholesterol probe ALOD4^27^ compared to ECs cultured in cholesterol-enriched media (10% FBS) (**Fig. 2a**). Furthermore, ECs cultured in cholesterol-depleted media showed blunted induction of VCAM-1 after LPS exposure (**Fig. 2a**). Similarly, limiting exogenous cholesterol supply by culturing HUVECs in 1% FBS for 24 h prior to and during a time course of LPS stimulation impaired VCAM-1 induction compared to culture in 10% FBS (both conditions included simvastatin to limit contributions of endogenous cholesterol synthesis) (**Extended Data Fig. 2a**). These data suggest that EC cholesterol availability influences the magnitude of VCAM-1 induction in response to inflammatory stimuli.

**Fig. 2.**
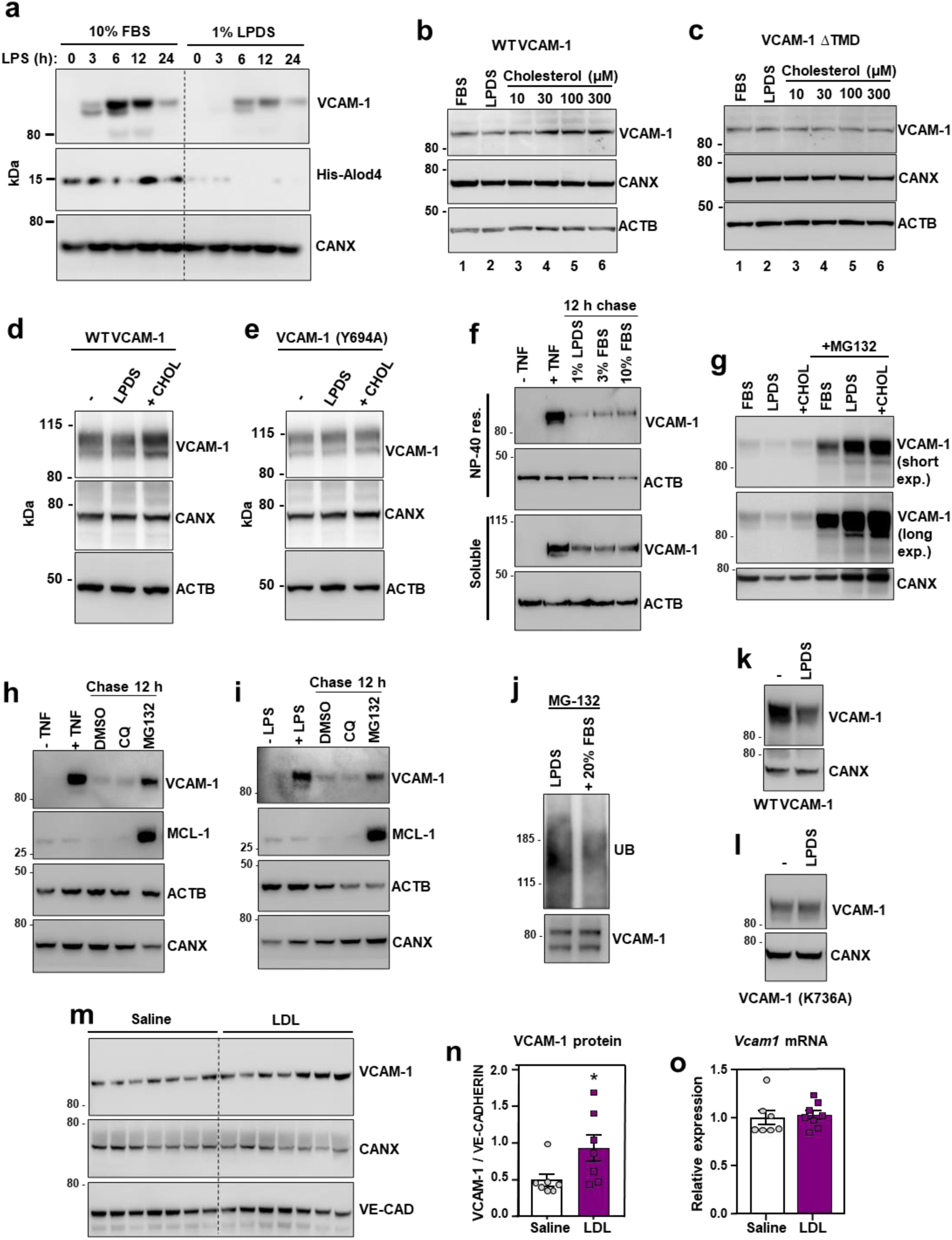
Cholesterol binding stabilizes VCAM-1. (a) Western blots for VCAM-1 and His-ALOD4 in HUVECs cultured in media containing either 10% FBS or 1% LPDS with simvastatin for 16 h before stimulation with LPS (100 ng/ml) for 0-24 h. (b and c) VCAM-1 western blots in cells stably overexpressing either human VCAM-1 or VCAM-1(ΔTMD) and cultured in media containing 10% FBS, 1% LPDS with simvastatin and mevalonate overnight, or LPDS with simvastatin and mevalonate overnight before addition of increasing concentrations of MβCD-cholesterol for 2 h. (d and e) VCAM-1 western blots in HUVECs stably expressing either FLAG-VCAM-1 or FLAG-VCAM-1(Y694A) cultured in media containing 10% FBS (-), 1% LPDS with simvastatin for 8 h, or 1% LPDS with simvastatin plus 100 µM MβCD-cholesterol for 2 h. (f) Western blots for VCAM-1 in HUVECs cultured in media containing 10% FBS and stimulated with TNFα for 12 h before being placed in media containing simvastatin and either 1% LPDS, 3% FBS or 10% FBS for a further 12 h. The top rows show the NP-40 resistant portion of cells while the bottom rows show the NP-40 soluble portion of cells. (g) VCAM-1 western blots showing the effects of proteasome inhibition with MG-132 in HUVECs stably overexpressing FLAG-VCAM-1 and cultured in 10% FBS, 1% LPDS with simvastatin overnight or LPDS with simvastatin plus 100 µM MβCD-cholesterol for 1 h. (h and i) HUVECs were treated with either LPS or TNFα for 36 h before being incubated with chloroquine (10 µM) or MG132 (10 µM) for a further 12 hours. VCAM-1 and MCL-1 (positive control for MG-132) were assessed by western blot. (j) HUVECs expressing FLAG-VCAM-1 were cultured in 1% LPDS with simvastatin overnight before being switched to media containing MG132 (10 µM) in 1% LPDS with simvastatin or 1% LPDS with simvastatin plus 20% FBS for 4 h. FLAG was immunoprecipitated before VCAM-1 and ubiquitin were assessed by western blot. (k and l) HUVECs expressing FLAG-VCAM-1 or FLAG-VCAM-1(K736A) were switched from media containing 10% FBS to media containing 1% LPDS with simvastatin for 12 h to assess their rate of degradation in cholesterol deplete conditions by western blotting. (m) Western blot for VCAM-1 in the lungs of male WT mice that received i.v. infusions of either saline or LDL for 6 h (n = 7 per group). (n) Quantification of VCAM-1 relative to Ve-cadherin measured by western blot in the lungs of WT mice after i.v. infusions of LDL or saline for 6 h. (o) mRNA levels of *Vcam1* relative to *36b4* in the lungs of male WT mice that received i.v. infusions of either saline or LDL for 6 h (n=7 saline and 8 LDL). Data are represented as mean ± SEM with individual mice represented by dots.

To specifically examine post-transcriptional effects of cholesterol on VCAM-1, we stably overexpressed the protein in ECs using a viral vector. HUVECs cultured in cholesterol-enriched media containing 10% FBS had higher VCAM-1 levels compared to ECs cultured in cholesterol-depleted media for 16 h (**Fig. 2b and Extended Data Fig. 2b**). Loading cholesterol-depleted cells with increasing concentrations of MβCD-cholesterol dose dependently increased levels of the WT VCAM-1 protein (**Fig. 2b**), but not a mutant version of VCAM-1 lacking its TMD (**Fig. 2c**). These data suggest that VCAM-1 must be anchored in a membrane to be regulated by cholesterol. Accordingly, the abundance of WT VCAM-1 was reduced when cells were switched to cholesterol-depleted media and was increased by re-introduction of cholesterol (**Fig. 2d**). However, the Y694A mutant had impaired regulation by cholesterol (**Fig. 2e**). Therefore, the CARC motif in the TMD of VCAM-1 is important for its regulation by cholesterol.

We next exposed ECs to tumor necrosis factor α (TNFα), a cytokine that activates ECs, for 12 h to induce endogenous VCAM-1. We then withdrew the TNFα and switched the cells to media containing varying amounts of cholesterol. Since many transmembrane proteins are detergent insoluble, we examined VCAM-1 in cellular fractions that were either soluble or resistant to the detergent NP-40. Culturing ECs in media containing 1% LPDS accelerated the degradation of endogenous VCAM-1 in the NP-40-resistant fractions compared to culture in 3% or 10% FBS (**Fig. 2f**). VCAM-1 levels were also higher in the detergent-soluble fraction in cells cultured in 10% FBS compared to 1% LPDS or 3% FBS (**Fig. 2f**). Given the apparent cholesterol-responsiveness of VCAM-1 in the NP-40-resistant portion of cells, we carried out further experiments on VCAM-1 in detergent-resistant domains. FLAG-VCAM-1 was depleted from detergent-resistant domains when ECs were cultured in media containing 1% LPDS and simvastatin compared to media containing 10% FBS and was robustly increased again after addition of MβCD-cholesterol or LDL (**Extended Data Fig. 2c**). The effects of cholesterol on VCAM-1 appeared to be more robust in the detergent-resistant compared to the detergent-soluble cell fraction. These observations imply that association with cholesterol makes VCAM-1 more resistant to solubilization by detergents.

The ability of cholesterol to regulate VCAM-1 abundance in ECs was ablated by the proteasome inhibitor MG132 (**Fig. 2g**). MG132, but not chloroquine, also slowed the degradation of endogenous VCAM-1 in LPDS after induction with TNFα or LPS (**Fig. 2h and 2i**). Immunoprecipitation experiments further showed that re-introduction of cholesterol in sterol-depleted HUVECs reduced the ubiquitination status of VCAM-1 in the presence of MG132 (**Fig. 2j**). Publicly available mass spec datasets suggested that lysine 736 (K736) in VCAM-1 may be ubiquitinated in human cells^28^. WT VCAM-1 protein was depleted after switching cells to LPDS for 12 h, but this effect was ablated by mutating K736 to alanine (K736A) (**Fig. 2k and 2l**). Together these data suggest that cholesterol binding to the TMD of VCAM-1 inhibits its degradation by limiting access of proteasomal machinery to lysine residues in the carboxy terminus tail.

To determine whether acute cholesterol delivery to the endothelium affects VCAM-1 abundance *in vivo*, we infused WT mice with freshly isolated LDL particles. Plasma total cholesterol levels were increased 6 h after intravenous (i.v.) infusions of LDL compared to control infusions of saline (**Extended Data Fig. 2d**). Furthermore, LDL infusions increased VCAM-1 protein levels in the lungs (**Fig. 2m and 2n**) without altering *Vcam1* mRNA levels (**Fig. 2o**). These data suggest that LDL acutely increases VCAM-1 in the lung by affecting its stability rather than its transcription.

### Inflammatory signals expand the accessible cholesterol pool to stabilize VCAM-1

We next examined PM cholesterol dynamics in ECs at baseline and after activation. Exposure to LPS or TNFα for 1 h increased ALOD4 binding to the surface of ECs, indicating increased cholesterol accessibility (**Fig. 3a**). It has been reported previously that TNFα and other proinflammatory agents activate PM-localized neutral sphingomyelinase (nSMase)^29–37^. We therefore hypothesized that SM hydrolysis induced by cytokines or LPS might liberate sequestered cholesterol for interactions with VCAM-1. Indeed, exposure to LPS or TNFα for 40 mins reduced ^3^H-SM levels in ECs relative to vehicle-treated control cells (**Fig. 3b**). Furthermore, incubating ECs with GW4869, a nSMase inhibitor, blunted the increase in ALOD4 binding induced by LPS or TNFα (**Fig. 3c**). Culturing ECs in the presence of methyl-β-cyclodextrin-cholesterol or exogenous nSMase served as controls for ALOD4 binding. (**Fig. 3c**).

**Fig. 3.**
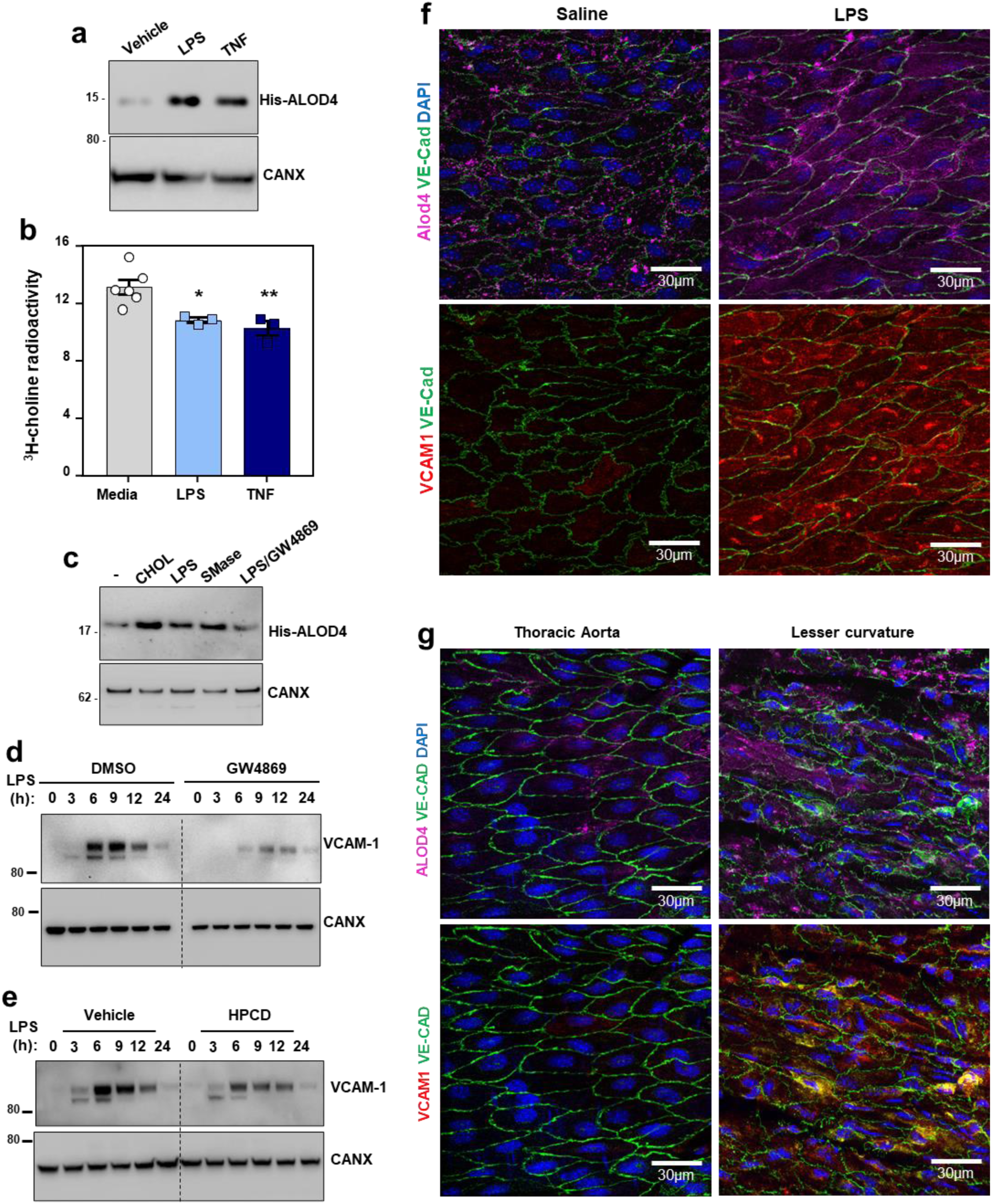
PM cholesterol accessibility increases during EC activation and influences the magnitude of VCAM-1 induction. (a) Western blot to assess His-ALOD4 binding to the surface of HUVECs exposed to either LPS (100 ng/ml) or TNFα (10 ng/ml) for 1 h. (b) [3H]-choline-labeled SM in HAECs after incubation with LPS (100 ng/ml) or TNFα (7.5 ng/ml) for 40 mins. (c) Western blot to assess His-ALOD4 binding to the surface of HAECs treated with MβCD-cholesterol (100 µM), LPS (100 ng/ml), bacterial nSMase (1 U/ml), or LPS co-incubated with the neutral sphingomyelinase inhibitor GW4869 (5 µM) for 1 h. (d) Western blot for VCAM-1 in HUVECs pre-treated with or without GW4869 (10 µM) for 30 mins before exposure to LPS for the indicated times. (e) VCAM-1 in HUVECs treated with LPS for 30 mins before being incubated with or without HPCD for 15 mins. After washing away the HPCD, cells were incubated with media containing 10% FBS for the indicated times. (f) ALOD4-647 binding to the thoracic aorta of female mice after i.p injections of either saline or LPS (60 µg per mouse) for 3 h. Samples were co-stained with VCAM-1 (red), Ve-cadherin (green) and DAPI (blue). Scale bar, 30 µm (g) ALOD4-647 binding to thoracic aorta or the aortic arch (lesser curvature) of female mice. Samples were co-stained with VCAM-1 (red), Ve-cadherin (green) and DAPI (blue). Scale bar, 30 µm. Data are represented as mean ± SEM.

Promoting SM hydrolysis with exogenous nSMase for 1 h increased the abundance of stably overexpressed VCAM-1 in ECs (**Extended Data Fig. 3a**). Conversely, blocking SM hydrolysis in response to LPS with GW4869 blunted the induction of endogenous VCAM-1 (**Fig. 3d**). Additionally, extracting the accessible PM cholesterol that appeared after LPS or TNFα exposure with HPCD lowered VCAM-1 protein levels over time (**Fig. 3e**). These data indicate that the newly accessible cholesterol pool released after EC activation is important for VCAM-1 induction.

To visualize accessible cholesterol dynamics in the vasculature, we perfused mice with fluorophore-conjugated ALOD4 (647 nm emission) through the left ventricle 3 h after intraperitoneal (i.p.) injection of saline or LPS. A separate set of mice was perfused with a cholesterol-binding mutant version of ALOD4 (G501A, T502A, T503A, L504A, Y505A, and P506A) to control for non-specific probe binding^27^. ALOD4-positive puncta were observed on the surface of the endothelium of mice that received control saline injections (**Fig. 3f**). Minimal signal was observed on the endothelium of mice that received infusions of the cholesterol-binding mutant version of ALOD4 (**Extended Data Fig. 3b**). LPS exposure increased ALOD4 binding to the endothelium, indicating a rise in cholesterol accessibility (**Fig. 3f**). Interestingly, the pattern of ALOD4 staining shifted from puncta at the periphery of ECs to a striated pattern across the surface of the cells (**Fig. 3f**). An increase in VCAM-1 protein levels accompanied the increase in ALOD4 binding to the endothelium after LPS injection (**Fig. 3f**). These data suggest that pro-inflammatory signals acutely increase cholesterol accessibility in the PM of ECs *in vivo*.

Previous work showed that VCAM-1 expression is higher in the lesser curvature of the aortic arch compared to the descending thoracic aorta^38^. We observed more ALOD4 binding to the lesser curvature of mouse aortas, where VCAM-1 was more highly expressed, compared to the descending aorta, where VCAM-1 expression was relatively low (**Fig. 3g**). This observation suggests that cholesterol availability correlates with VCAM-1 abundance on the endothelium *in vivo*.

We next developed a protocol to assess ALOD4 binding to fixed mouse aortas *ex vivo*. Using this protocol, we observed that accessible cholesterol levels in the aortic endothelium of atherosclerotic LDLR knockout mice were higher than those in the aortas of normocholesterolemic mice fed a regular chow diet (**Extended Data Fig. 3c**). These data suggest that circulating cholesterol concentrations influence the size of the accessible cholesterol pool on the endothelium.

### The cholesterol transporter Aster-A is engaged during EC activation

Our experiments thus far suggested that the PM accessible cholesterol pool acutely increases during EC activation to allow cholesterol interactions with VCAM-1. Since Aster proteins regulate the size of the PM accessible cholesterol pool in various cell types^9,10,39^, we investigated their role during EC activation. Primary ECs isolated from the livers of WT mice expressed high levels of Aster-A, with comparatively low levels of Aster-B and -C (**Fig. 4a**). Cultured primary human aortic endothelial cells (HAECs) also predominantly expressed Aster-A (**Extended Data Fig. 4a**). Analysis of published single-cell sequencing datasets confirmed high Aster-A expression and low Aster-B and -C expression in mouse ECs from different tissue beds^40^.

**Fig. 4.**
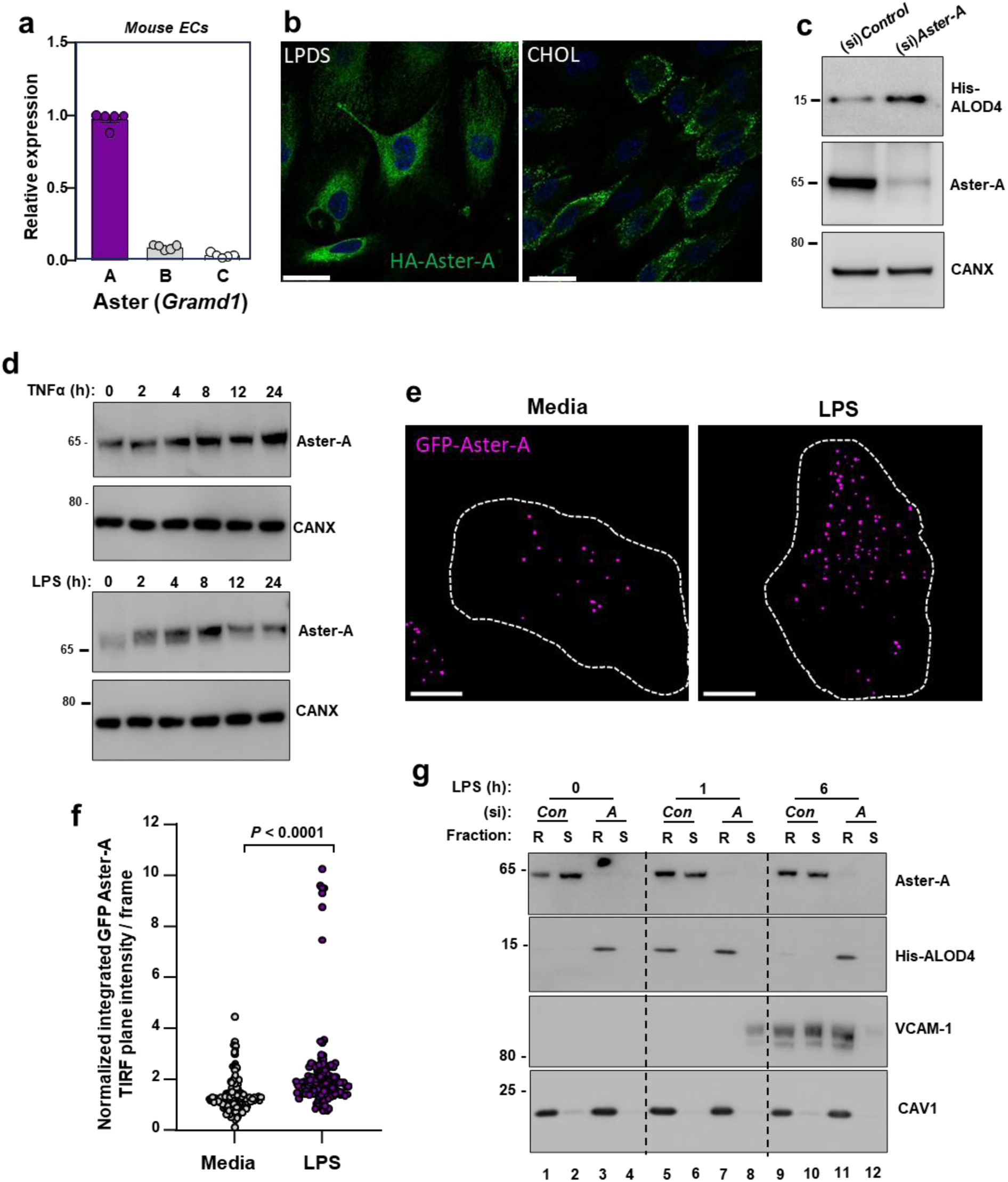
Aster-A regulates accessible cholesterol in ECs after activation. (a) *Gramd1a* (Aster-A), *Gramd1b* (Aster-B) and *Gramd1c* (Aster-C) expression relative to *36b4* in primary mouse hepatic ECs. (b) Confocal microscopy of HAECs stably expressing HA-Aster-A and cultured in LPDS or LPDS with MβCD-cholesterol (100 µM) for 1 h. Scale bar, 23 µm. (c) His-ALOD4 binding to the surface of HAECs treated with (si)Control or (si)Aster-A and cultured in 10% FBS. (d) Endogenous Aster-A protein levels in HUVECs exposed to TNFα (10 ng/ml; top) or LPS (100 ng/ml; bottom) for the indicated times. (e) TIRF microscopy of HAECs stably expressing EGFP-Aster-A and cultured in fresh complete medium (10% FBS) or fresh complete medium plus LPS (100 ng/ml) for 60 mins. Pseudo-colored dots indicate GFP-Aster-A intensity in the TIRF plane (within 100 nm of the PM). Dashed lines indicate cell boundaries. Scale 10 µm. (f) Quantification of GFP-Aster-A in the TIRF plane +/- LPS. Values represent normalized integrated intensities at 40-70 min after changing to fresh media +/- LPS. Control n = 154 frames from 52 cells, LPS n = 140 frames from 53 cells from two independent experiments. (g) Western blots of HUVECs treated with (si)Control or (si)Aster-A and exposed to LPS for the indicated times before incubation with ALOD4. Cells were subsequently fractionated into Trition-X100 detergent soluble or detergent resistant domains. Data are represented as mean ± SEM.

HA-tagged Aster-A was distributed throughout the ER in HAECs cultured in media containing LPDS with simvastatin, but was recruited to the PM in response to 1 h cholesterol loading (**Fig. 4b**). Aster-A depletion with a small interfering (si)RNA increased ALOD4 binding to the surface of HAECs (**Fig. 4c**), consistent with a major role for Aster-A in regulating PM accessible cholesterol levels in ECs. *GRAMD1A* mRNA levels were not changed in response to TNFα exposure of HAECs (**Extended Data Fig. 4b**). Notably, however, the endogenous Aster-A protein increased with time after TNFα (**Extended Data Fig. 4c**). Aster-A protein levels also increased in primary HUVECs after exposure to TNFα or LPS (**Fig. 4d**). Furthermore, HA-tagged Aster-A expressed from a viral vector increased with time after EC activation (**Extended Data Fig. 4d**). Together these data suggest that the cholesterol transporter Aster-A undergoes post-transcriptional stabilization in response to pro-inflammatory signals in ECs. Regulation of Aster-A by cytokines and LPS is consistent with a role for nonvesicular cholesterol transport during EC activation.

### Asters remove accessible cholesterol from EC PMs after activation

The acute rise in accessible cholesterol in response to inflammatory signals in ECs would be predicted to recruit Aster to the PM to move cholesterol to the ER. Indeed, total internal reflection (TIRF) microscopy showed that EGFP-Aster-A was recruited to the TIRF plane (indicating proximity to the PM) in response to LPS exposure (**Fig. 4e and 4f**). GFP-Aster-A was enriched in the TIRF plane 40-70 mins after LPS exposure compared to baseline media conditions (**Fig. 4f**). These data indicate the EC activation results in Aster recruitment to the PM.

To determine whether Aster-A plays a role in accessible cholesterol transport downstream of EC activation, cells were incubated with ALOD4 at various times after LPS exposure before being fractionated into detergent-resistant and detergent-soluble domains. PM-bound ALOD4 specifically partitioned into detergent-resistant membrane domains, defined by the presence of CAV1 (**Fig. 4g**). Control ECs had low accessible cholesterol levels at baseline (Lane 1), a rise in accessible cholesterol after 1 h LPS exposure (Lane 5), and a return to baseline by 6 h (Lane 9; **Fig. 4g**). Aster-A deficient ECs had higher ALOD4 binding at baseline (Lane 3) and 1 h after LPS exposure (Lane 7) relative to control cells. Furthermore, in contrast to control cells, Aster-A deficient cells failed to remove the PM accessible cholesterol by 6 h after LPS exposure (Lane 11; **Fig. 4g**). Endogenous Aster-A protein partitioned predominantly into the detergent soluble domains at baseline in control cells (Lane 2), was recruited to the ALOD4-positive detergent-resistant domains in response to 1 h LPS exposure (Lane 5), and largely returned to the detergent-soluble fraction after 6 h (Lane 10) when PM accessible cholesterol had been depleted. Consistent with the accessible cholesterol pool playing a role in VCAM-1 stability, the ALOD4-positive detergent-resistant domains of Aster-deficient ECs contained higher levels of VCAM-1 6 h after LPS exposure compared to control cells (Lane 11; **Fig. 4g**). Therefore, EC activation promotes Aster translocation to accessible cholesterol-enriched regions of the PM to move cholesterol to the ER.

### Aster-A regulates VCAM-1 stability in ECs

Loss of Aster-A increased VCAM-1 in HAECs after activation with LPS (**Extended Data Fig. 5a**), and VCAM-1 protein was strikingly higher in detergent-resistant fractions of Aster-deficient ECs 6 h after LPS exposure (**Fig. 5a**). Interestingly, Aster-A deficiency also modestly increased the abundance of *VCAM1* transcripts 4 h after LPS compared to control cells (**Fig. 5b**). We therefore hypothesized that PM cholesterol plays a dual role during EC activation: first by modulating the PM-derived signals that determine the magnitude of *VCAM1* transcriptional induction, and then directly binding to the translated products at the PM. This hypothesis was based in part on previous observations that excess PM cholesterol enhances the recruitment of the pro-inflammatory signaling adapters TRAF6 and MYD88 to detergent resistant membrane domains after TLR4 agonism in macrophages, which amplifies downstream signaling^41^. We found that loss of Aster-A increased ALOD4 binding to detergent-resistant domains of HUVECs and caused more MYD88 and TRAF6 to localize to ALOD4-positive domains 15 mins after LPS exposure compared to control cells (**Fig. 5c**; lane 2 compared to lane 6). Loss of Aster-A also increased the phosphorylation of p44 and p42 MAPK after LPS exposure (**Fig. 5d**), consistent with amplified signaling downstream of TRAF6. Loading ECs with cholesterol was sufficient to promote TRAF6 and MYD88 recruitment to CAV1/FLOT1-positive detergent-resistant domains (**Extended Data Fig. 5b**; lane 2 compared to lane 6), suggesting that the effects of Aster deficiency on TRAF6/MYD88 were mediated by cholesterol. Loading cells with cholesterol also promoted Aster-A translocation from detergent-soluble domains to detergent-resistant domains (**Extended Data Fig. 5b**; lane 2 compared to lane 6). Additionally, loading cholesterol-depleted ECs with FBS-cholesterol for 4 h increased mRNA levels of *VCAM1* and a panel of other NF-KB targets while suppressing SREBP-2 targets (**Extended Data Fig. 5c**). Following exposure to LPS, the magnitude of *VCAM1* induction was higher when ECs were cultured in cholesterol-enriched (10% FBS) compared to cholesterol-depleted (1% LPDS with simvastatin) conditions (**Extended Data Fig. 5d**). Thus, accessible cholesterol accumulation on EC PMs promotes the transcription of *VCAM1* by amplifying TRAF6/MYD88 signaling.

**Fig. 5.**
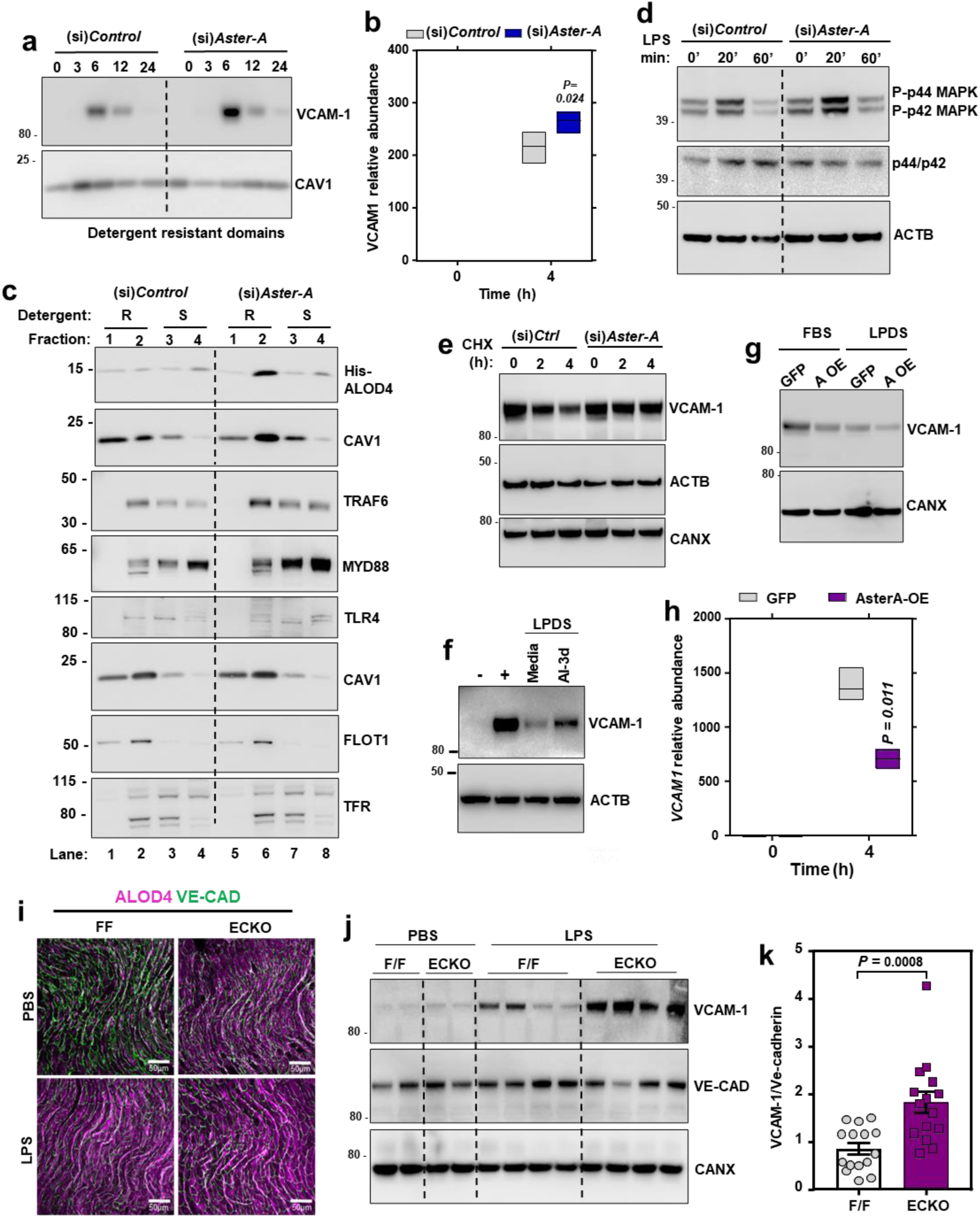
Nonvesicular cholesterol transport regulates VCAM-1 stability *in vivo*. (a) Western blot for VCAM-1 in detergent resistant domains of immortalized HAECs treated with (si)Control or (si)Aster-A and exposed to LPS (100 ng/ml) for the indicated times. Cells were cultured in media containing 5 % FBS and simvastatin for 16 h before LPS exposure. (b) *VCAM1* mRNA levels relative to *36B4* in HUVECs exposed to LPS for the indicated times. Center line, mean; box limits, upper and lower values. (c) Western blots in HUVECs treated with (si)Control or (si)Aster-A and exposed to LPS for 15 mins before fractionation into detergent resistant (fractions 1 or 2) or detergent soluble domains (fractions 3 or 4). Cells were cultured in media containing 5% FBS with simvastatin overnight before exposure to LPS. Separate dishes were used to assess -His-ALOD4 binding (top 2 rows) and TRAF6/MYD88 localization (bottom 6 rows). (d) Western blots for p-ERK or total ERK in HUVECs treated with (si)Control or (si)Aster-A and exposed to LPS (100 ng/ml) for the indicated times. (e) Cycloheximide chase of FLAG-VCAM-1 in HUVECs treated with (si)Control or (si)Aster-A. (f) Western blots for VCAM-1 HUVECs stimulated with LPS for 12 h before being placed in media containing LPDS and simvastatin with or without AI-3d (2.5 µM) for a further 12 h. (g) VCAM-1 protein levels in HAECs stably overexpressing Aster-A or GFP and cultured in either 10% FBS or 1% LPDS with simvastatin before exposure to LPS (100 ng/ml) for 8 h. (h) *VCAM1* mRNA levels relative to *36B4* in HAECs stably overexpressing HA-Aster-A or GFP and exposed to LPS for the indicated times. Center line, mean; box limits, upper and lower values. (i) ALOD4-647 binding to aortas of male F/F and ECKO mice injected with either saline or LPS (60 µg per mouse) for 3 h. Samples were co-stained with Ve-cadherin (green). Scale bar, 50 µm. (j) Western blots for VCAM-1 in the hearts of male F/F and ECKO mice 3 h after i.p. injections of saline or LPS (60 µg/mouse). (k) Quantification of VCAM-1 relative to Ve-cadherin measured by western blot in the hearts of male F/F and ECKO after i.p. injections of LPS (60 µg/mouse) for 3 h. n = 15 mice per group. Data is from 3 independent experiments. Data are represented as mean ± SEM.

To directly examine the effects of cellular cholesterol transport on VCAM-1 stability, we manipulated Aster function in cells that stably overexpressed VCAM-1. Loss of Aster-A increased the stability of FLAG-VCAM-1 during a cycloheximide chase (**Fig. 5e**). Additionally, Aster inhibition with the small molecule AI-3d^42^ slowed the rate of degradation of endogenous VCAM-1 in ECs after induction with LPS (**Fig. 5f**).

We next overexpressed Aster-A in HAECs to promote accessible cholesterol movement from the PM to the ER. In HAECs expressing GFP treated with LPS, VCAM-1 abundance was lower when cells were cultured in cholesterol-depleted (LPDS with simvastatin) compared to cholesterol-enriched media (10% FBS; **Fig. 5g**). HAECs over-expressing Aster-A cultured under similar conditions showed further reduced VCAM-1 protein levels. Additionally, the magnitude of *VCAM1* transcriptional induction by LPS was lower in cells overexpressing Aster-A compared to control cells overexpressing GFP (**Fig. 5h**). These data further implicate PM cholesterol availability and the flux of cholesterol from the PM to intracellular membranes as regulators of VCAM-1 abundance in ECs.

### Impairing cholesterol transport from the PM increases VCAM-1 in ECs in vivo

To study PM accessible cholesterol on the endothelium *in vivo*, we crossed Aster-A-floxed mice (F/F) with mice expressing a *Cdh5*-Cre inducible with tamoxifen^43^. *Gramd1a* mRNA (**Extended Data Fig. 5e**) and Aster-A protein levels (**Extended Data Fig. 5f**) were undetectable in primary ECs isolated from Aster-A^F/F^/*Cdh5*-Cre^-/+^ (ECKO) mice compared to F/F control mice. Liver tissue from ECKO mice had comparable *Gramd1a* mRNA levels to F/F controls (**Extended Data Fig. 5g**), consistent with the EC specificity of the *Cdh5*-Cre system. Importantly, ECKO mice had higher ALOD4 binding to the aortic endothelium compared to F/F control mice 3 h after i.p. saline injection (**Fig. 5i**). Injection of LPS for 3 h increased ALOD4 binding to the endothelium of F/F control mice relative to saline injection and increased it further in ECKO mice (**Fig. 5i**). VCAM-1 protein was low in the hearts of F/F and ECKO mice after control saline injections (**Fig. 5j**), reflecting the low basal expression in endocardial ECs. However, VCAM-1 protein levels were over 2-fold higher in the hearts of ECKO mice compared to F/F controls 3 h after i.p. LPS (**Fig. 5j** and **5k**). These data indicate that Aster-A nonvesicular cholesterol transport regulates accessible PM cholesterol and the magnitude of VCAM-1 induction *in vivo*.

### Cholesterol transport from the PM to intracellular membranes of ECs maintains lung immune homeostasis

To determine whether Aster-A participates in lipoprotein-cholesterol movement across the endothelium into tissues, we conducted tracer studies with freshly isolated HDL particles labeled with [^14^C]-cholesterol. The uptake of i.v. administered [^14^C]-cholesterol-HDL into the lungs of ECKO was dramatically lower compared to F/F mice, while uptake into most other tissues was similar between groups (**Fig. 6a**). This observation suggested that the pulmonary endothelium might be particularly affected by loss of Aster-A. Western blots showed higher VCAM-1 protein levels in the lungs of ECKO mice compared to F/F controls after both acute (3 weeks) (**Extended Data Fig. 6a**) and chronic (1 year) (**Fig. 6b and 6c**) Cre induction. Immuno-fluorescence microscopy also indicated that ECKO mice had higher VCAM-1 throughout the lungs compared to F/F controls (**Fig. 6d**). Co-staining with ERG, an EC-specific nuclear marker, revealed that most VCAM-1 in the lungs was associated with ERG-positive cells (**Fig. 6d**). H&E-stained lung sections showed immune cells around vessels in the lungs of ECKO mice (**Fig. 6e**), suggesting that higher basal VCAM-1 abundance in the lung was associated with immune cell recruitment to the pulmonary endothelium. We also found more CD45-positive immune cells around Lyve1-positive vessels in the lungs of ECKO mice (**Fig. 6f**). Therefore, nonvesicular cholesterol transport in ECs influences adhesion molecule abundance, lung immune homeostasis, and immune cell recruitment to the endothelium *in vivo*.

**Fig. 6.**
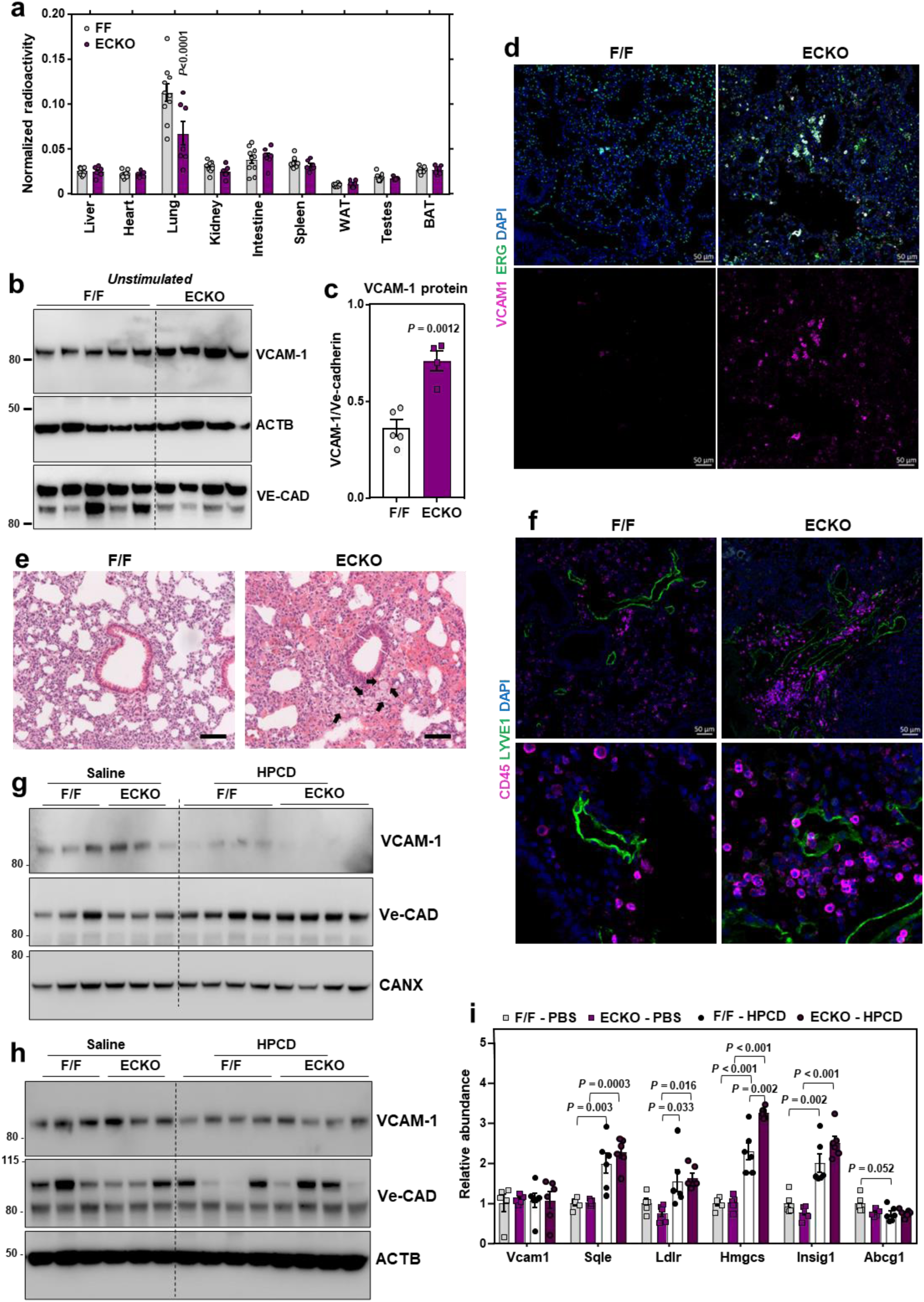
HPCD infusions lower VCAM-1 in response to LPS *in vivo*. (a) Tissue [^3^H]-cholesterol radioactivity normalized to tissue weight in male F/F and ECKO mice 72 h after i.v. administration of [^3^H]-cholesterol-HDL. Samples analyzed by two-way ANOVA with genotype and tissue as independent variables. *P* tissue < 0.0001; *P* genotype < 0.0061; *P* interaction < 0.0001. n = 10 F/F and 7 ECKO. (b) Western blots for VCAM-1 in the lungs of female F/F and ECKO mice 1 year after Cre induction. n = 5 F/F and 4 ECKO. (c) Quantification of VCAM-1 relative to Ve-cadherin measured by western blot as shown in Fig. 7B. (d) Immunofluorescence microscopy of VCAM-1 (pink), ERG (green) and DAPI (blue) in the lungs of female F/F and ECKO mice 3 weeks after Cre induction. Scale bar, 50 µm. (e) H & E staining in the lungs of female F/F and ECKO mice 3 weeks after Cre induction. Arrows indicate immune cells around vessels. Scale bar, 100 µm. (f) CD45-positive immune cells (purple) co-stained with the lymphatic vessel marker LYVE1 (green) and DAPI (blue) in the lungs of female F/F and ECKO mice 3 weeks after Cre induction. Scale bar, 50 µm. (g and h) VCAM-1 in the hearts and lungs of male F/F and ECKO mice injected with LPS for 20 mins before receiving i.v infusions of saline or HPCD. Tissues were collected 3 h after LPS injections. n = 5 F/F + saline, 5 ECKO + saline, 6 F/F + HPCD and 6 ECKO + saline. (i) qPCR in the lungs of male F/F and ECKO mice injected with LPS for 20 mins before receiving i.v infusions of saline or HPCD. Tissues were collected 3 h after LPS injections. n = 5 F/F + saline, 5 ECKO + saline, 6 F/F + HPCD and 6 ECKO + saline. Data are represented as mean ± SEM with individual mice noted as dots.

### Acute HPCD infusions reduce VCAM-1 induction in response to LPS

Our data thus far suggested that direct interaction between VCAM-1 and accessible cholesterol stabilizes the protein on the EC surface. HPCD extracts cholesterol from membranes and has U.S. Food and Drug Administration (FDA) approval for use in humans, primarily for its ability to solubilize hydrophobic compounds^44^. We hypothesized that extraction of cholesterol from the surface of ECs with i.v. infusions of HPCD during an infection might destabilize VCAM-1. To test this hypothesis, F/F and ECKO mice were injected with LPS for 20 min before receiving i.v. infusions of HPCD. Two hours and 40 min later, tissues were collected for assessment of VCAM-1. HPCD infusions lowered total plasma cholesterol concentrations in both F/F and ECKO mice (**Extended Data Fig. 6b**), likely due to cholesterol solubilization and removal by the kidneys^45^. HPCD infusions also resulted in a dramatic decrease in cardiac and pulmonary VCAM-1 protein levels in both F/F and ECKO mice compared to saline infusions (**Fig. 6g and 6h**). Heart and lung qPCR analysis showed no difference in *Vcam1* transcripts in the presence or absence of HPCD, suggesting that HPCD altered VCAM1 stability rather than production in these tissues (**Fig. 6i and Extended Data Fig. 6c**). HPCD infusions did not lower protein levels of Ve-cadherin, another cell surface-localized adhesion molecule (**Fig. 6g and 6h**). SREBP-2 target genes (*Sqle*, *Hmgcs*, *Insig1*) were increased in response to HPCD administration in the lungs of F/F and ECKO mice, consistent with cholesterol extraction from cell membranes (**Fig. 6i**). Modest induction of SREBP-2 targets after HPCD was also observed in the heart (**Extended Data Fig. 6c**). Collectively, these data show that direct cholesterol stabilization of VCAM-1 protein is an important modulator of EC function in physiology and pathophysiology.

## Discussion

Activated ECs display VCAM-1 on their PM to promote leukocyte recruitment to injured vessels^3,4^. Monocyte adherence to vascular ECs is one of the earliest changes observed after initiation of a high-cholesterol atherogenic diet^14–16^. High-cholesterol diet feeding rapidly increases VCAM-1 protein levels on the vasculature, before the appearance of monocytes in the intima^46^. Additionally, VCAM-1 has been localized to atherosclerosis-prone regions of arteries and is abundant in ECs overlying early foam cell-enriched lesions^38,47–49^. Mice deficient in VCAM-1 are protected against atherosclerosis when crossed onto a hypercholesterolemic background^50^. Our data shows that hypercholesterolemia increases the pool of accessible cholesterol on the surface of the endothelium in isolated ECs and *in vivo*, and that this pool of cholesterol acts to stabilize VCAM-1 by direct interactions. Pro-inflammatory stimuli including LPS also acutely increase the accessible cholesterol content of EC PMs. These observations suggest that ECs increase cell surface cholesterol availability during EC activation as a mechanism to accommodate more VCAM-1 in the PM, and that this mechanism could be subject to maladaptation during chronic hypercholesterolemia. Therefore, strategies that disturb cholesterol-VCAM-1 interactions might reduce vascular adhesiveness to immune cells in the setting of hypercholesterolemia or other pro-inflammatory events.

Identifying biologically important interactions between lipids and membrane proteins has been challenging, because both lipids and TMDs are poorly soluble and hydrophobic^51^. Advances in click chemistry and chemoproteomics have enabled the enrichment and identification of lipid-interacting proteins in living cells with proteome-wide coverage. We synthesized a new cholesterol mimetic probe, NBII-165, and used it to probe protein-sterol interactions in ECs. Prior studies in HeLa cells used sterol-mimetic probes with a diazirine group on the B-ring of the steroid core ^52,53^. It is likely that VCAM-1 was not identified as a cholesterol binding protein in these previous studies because VCAM-1 expression is largely restricted to activated ECs. While we focused on characterizing cholesterol interactions with VCAM-1 in this manuscript, there are other proteins on the list of interactors not previously known to bind cholesterol. Further studies of these proteins will likely provide insight into cholesterol-regulated processes in ECs.

The ability of ECs to rapidly alter their proteome upon activation is a key feature of their biology. Quiescent ECs express low levels of VCAM-1, but its transcription is robustly induced in response to various activating stimuli^1^. Our experiments showed that changes in the lipid composition of the PM play a role in facilitating the rapid increase in VCAM-1 protein levels upon EC activation. Cholesterol binding at the PM stabilizes VCAM-1, preventing its degradation by the proteosome. Mutating Y694 in the CARC motif of VCAM-1 diminished the ability of cholesterol to stabilize the molecule, indicating that cholesterol acts directly on the protein, rather than indirectly by altering membrane properties like thickness or fluidity. Proteasome inhibition also ablated the ability of cholesterol to regulate VCAM-1, consistent with a model in which cholesterol binding to the TMD of VCAM-1 results in a conformational change to the carboxy-terminus tail of the molecule to limit access to ubiquitin ligases. Further structural studies are required to confirm this idea. Nevertheless, acutely depleting cholesterol from the endothelium decreases VCAM-1, suggesting that cholesterol-VCAM-1 interactions are functionally important in physiology.

Aster proteins are known to participate in cholesterol transfer from the PM to the ER in cells specialized for steroid metabolism^9,11–13,54^. However, the ability of nonvesicular cholesterol transport to modulate the function of other cell types through changing PM lipid composition has not been explored. Unexpectedly, we found that Aster-A was recruited to the PM upon EC activation and was stabilized in response to pro-inflammatory signals. These observations implied a previously unrecognized role for intracellular sterol transport in vascular homeostasis. We further revealed that Asters gate the interaction between accessible PM cholesterol and the integral PM protein VCAM-1 during EC activation. Loss of Aster-A increases VCAM-1 abundance in response to EC activation, while Aster-A overexpression potently suppresses VCAM-1 induction. Therefore, PM lipid remodeling by the cholesterol transporter Aster-A calibrates the magnitude of VCAM-1 induction and facilitates the transition of resting ECs to an activated state.

The PM is a critical site for lipid second messenger production across all kingdoms of life. PM phospholipids and sphingolipids undergo enzymatic hydrolysis in response to acute stress to produce the signaling molecules ceramide, diacylglycerol, and inositol triphosphate^55^. On the endothelium, neutral sphingomyelinase activity is stimulated by various pro-inflammatory agents, including TNFα, IL1β, and oxidized LDL, producing ceramide and phosphorylcholine from SM^29,30,33,35,37^. Ceramide produced after SM hydrolysis acts as a second messenger to amplify pro-inflammatory signaling^55^. Our data suggest that accessible cholesterol released following disturbances to SM/cholesterol complexes could also have second messenger-like properties. We found that increased PM accessible cholesterol amplifies signaling downstream of the pro-inflammatory adapters MYD88 and TRAF6. Following its transcriptional induction, PM cholesterol directly binds to the VCAM-1 protein to promote its stability. Therefore, PM accessible cholesterol appears to act as an overarching signal that calibrates the magnitude of VCAM-1 induction in ECs. The observation that accessible cholesterol levels remain inappropriately elevated after EC activation in the absence of Aster-A suggests a physiological role for nonvesicular sterol transport in resetting PM lipid composition and dampening inflammatory signaling following EC activation.

In conclusion, these results demonstrate that dynamic changes in PM accessible cholesterol content modify EC adhesiveness. As a critical vascular adhesion molecule, VCAM-1 is involved in the development of many immune-mediated disorders including atherosclerosis, sepsis, and cancer^1^. The discovery that direct cholesterol-VCAM-1 interactions are required to stabilize the protein *in vivo* suggest that these interactions could be targeted to reduce VCAM-1 expression in pathological settings.

---

**Extended Data Fig. 1.**
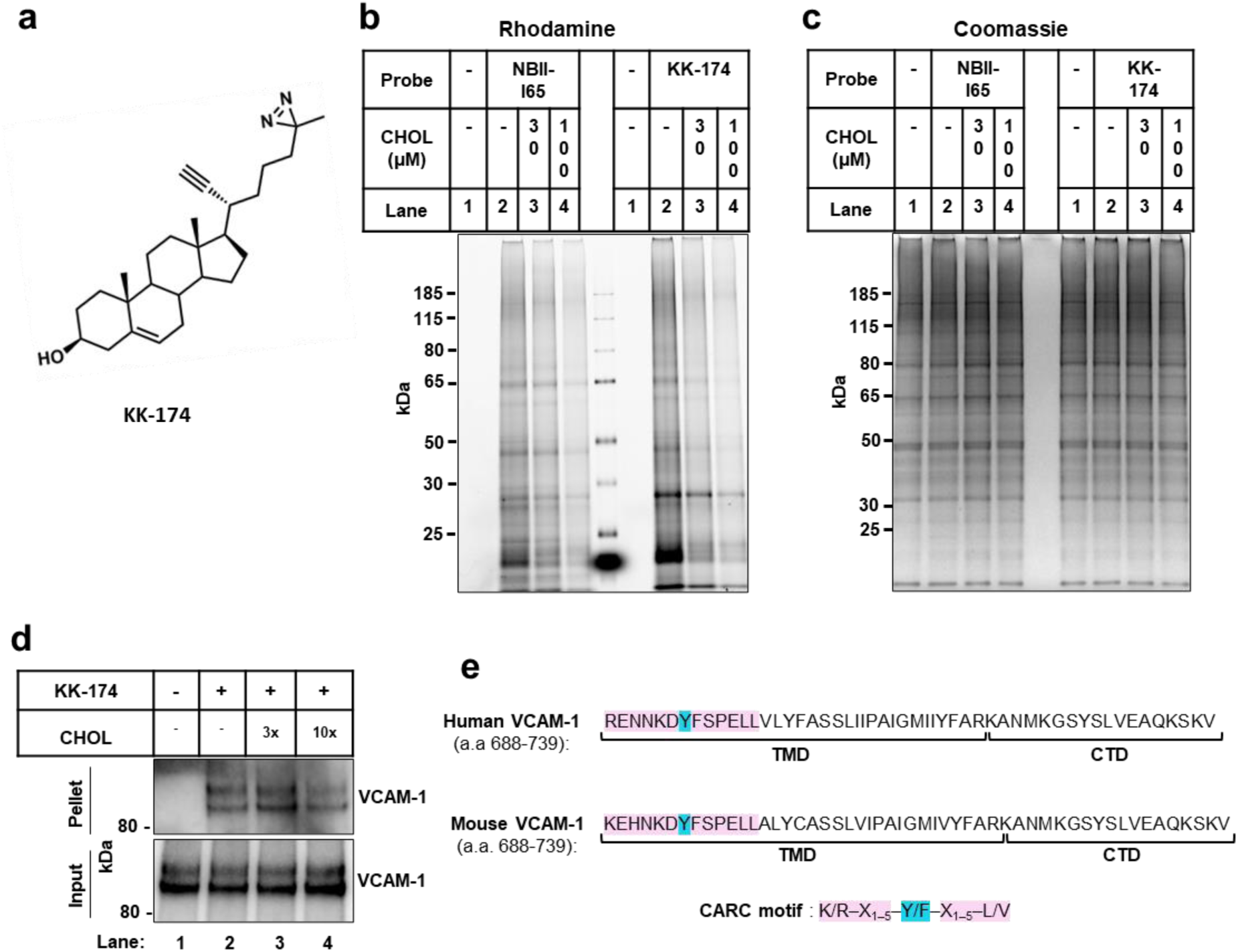
Cholesterol directly binds to VCAM-1. (a) Structure of the KK-174 probe. (b and c) The first lane in each condition represents cells with no cholesterol probe added before UV exposure. The remaining lanes represent cells loaded with 10 µM cholesterol-mimetic probe alone or with increasing concentrations of MβCD-cholesterol (30 µM or 100 µM) for 1 h before UV crosslinking. After UV exposure, cells were lysed, a rhodamine-azide tag was conjugated via click chemistry onto probe-bound samples, and cellular proteins were separated by SDS-PAGE. (b) Probe bound samples were visualized via the florescent rhodamine signal. (c) Coomassie staining of total cellular proteins from (b). (d) Competition assay showing that cholesterol competes with KK-174 for binding to VCAM-1 in HUVECs stably overexpressing human VCAM-1. Input shows VCAM-1 detected in whole cell lysates prior to immunoprecipitation and pellet shows VCAM-1 detected after streptavidin immunoprecipitation of probe bound proteins. (e) Amino acids 688-739 (corresponding to the TMD and CTD) in human and mouse VCAM-1 with the CARC motif in the TMD highlighted in pink and the central tyrosine (Y) residue that was mutated in Fig. 1F highlighted in blue.

**Extended Data Fig. 2.**
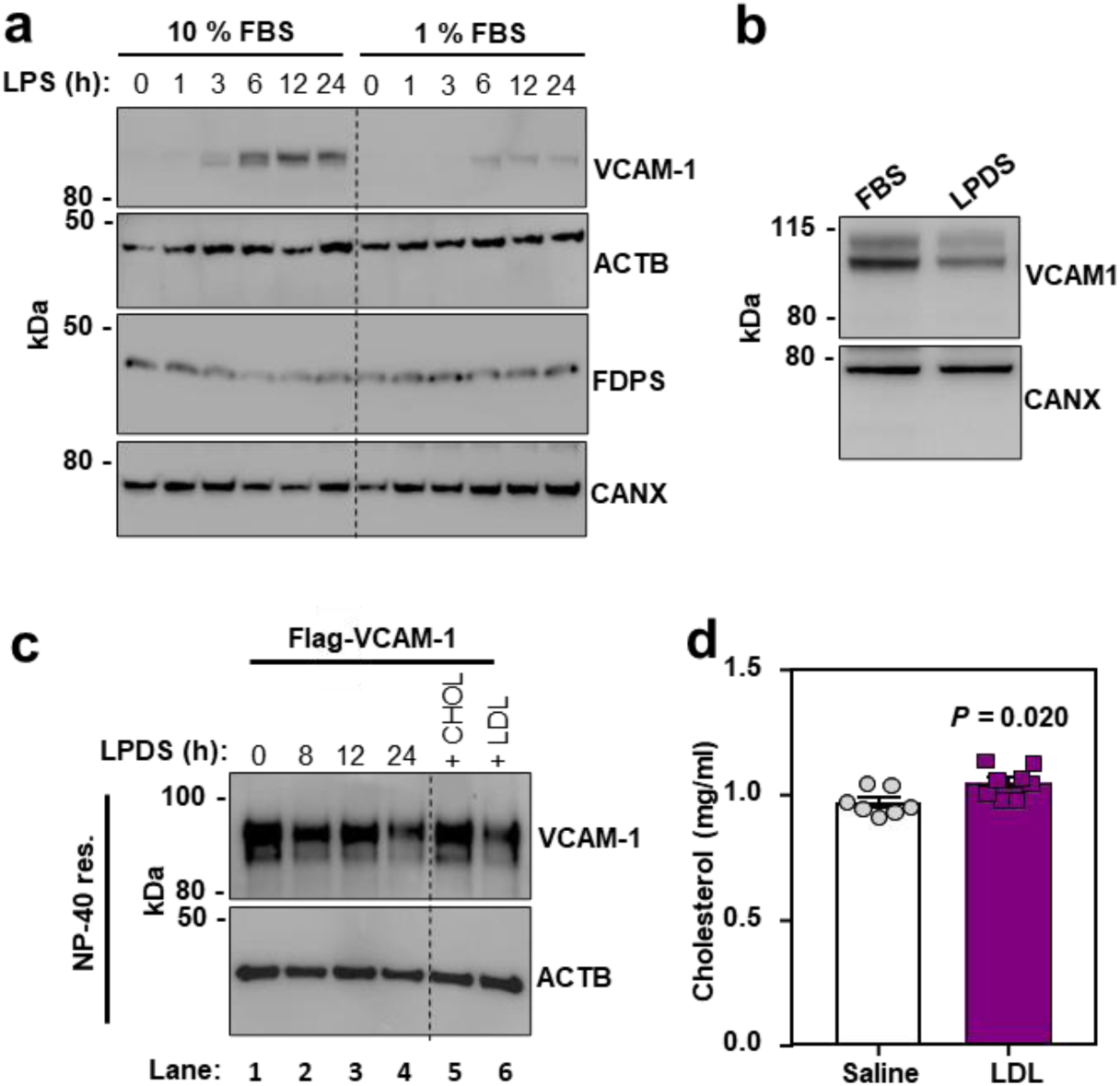
Cholesterol binding stabilizes VCAM-1. (a) Western blot for VCAM-1 in HUVECs cultured in media containing either 10% FBS with simvastatin or 1% FBS with simvastatin for 24 h before stimulation with LPS (100 ng/ml) for 0-24 h. (b) VCAM-1 western blots in cells stably overexpressing human VCAM-1 and cultured in media containing either 10% FBS or 1% LPDS with simvastatin for 16 h. (c) VCAM-1 western blots in the NP-40 resistant fraction of HUVECs stably overexpressing FLAG-VCAM-1 and cultured in media containing 1% LPDS with simvastatin for the indicated times. In the last two lanes, 100 µM MβCD-cholesterol or LDL (50 ug/ml) was added for the last 2 or 4 h of the 24 h LPDS chase, respectively. (d) Total plasma cholesterol in male WT mice 6 h after receiving i.v. infusions of either saline or LDL (n = 7 saline and 8 LDL). Data are represented as mean ± SEM with individual mice noted as dots.

**Extended Data Fig. 3.**
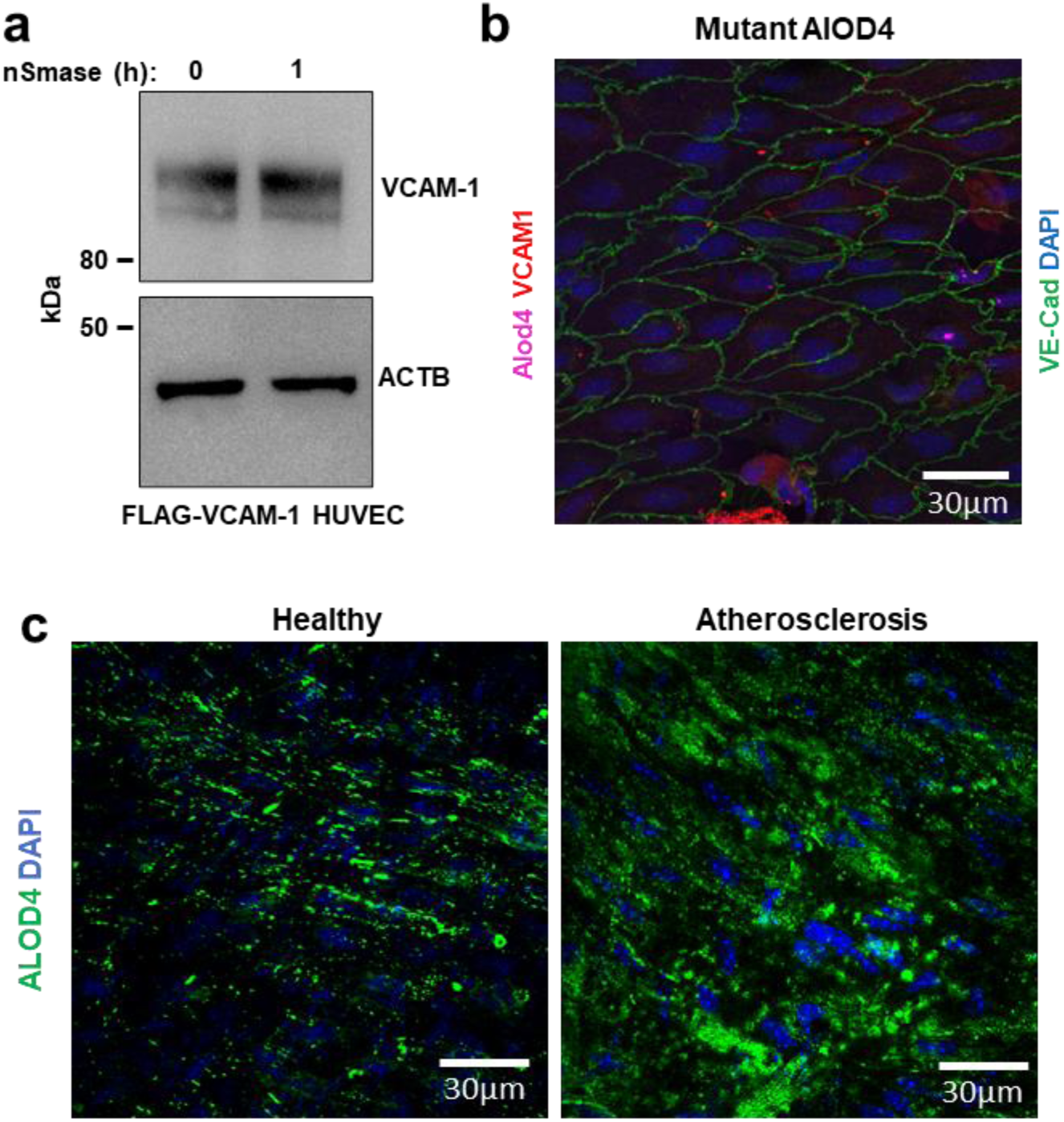
PM cholesterol accessibility increases during EC activation and hypercholesterolemia. (a) HUVECs stably expressing FLAG-VCAM-1 and cultured in media containing 1% LPDS with simvastatin overnight were treated with or without neutral sphingomyelinase for 2 h before immunoblotting for VCAM-1. (b) *En face* imaging of aortas from female mice that had been perfused with cholesterol-binding mutant versions ALOD4-647. Samples were co-stained with VCAM-1 (red), Ve-cadherin (green), and DAPI (blue). Scale bar, 30 µm. (c) ALOD4-488 binding to *en face* aortas of either male WT mice fed a chow diet or LDLR knockout mice that had been fed a Western diet for 20 weeks to induce atherosclerosis. Samples were co-stained with DAPI (blue). Scale bar, 30 µm.

**Extended Data Fig. 4.**
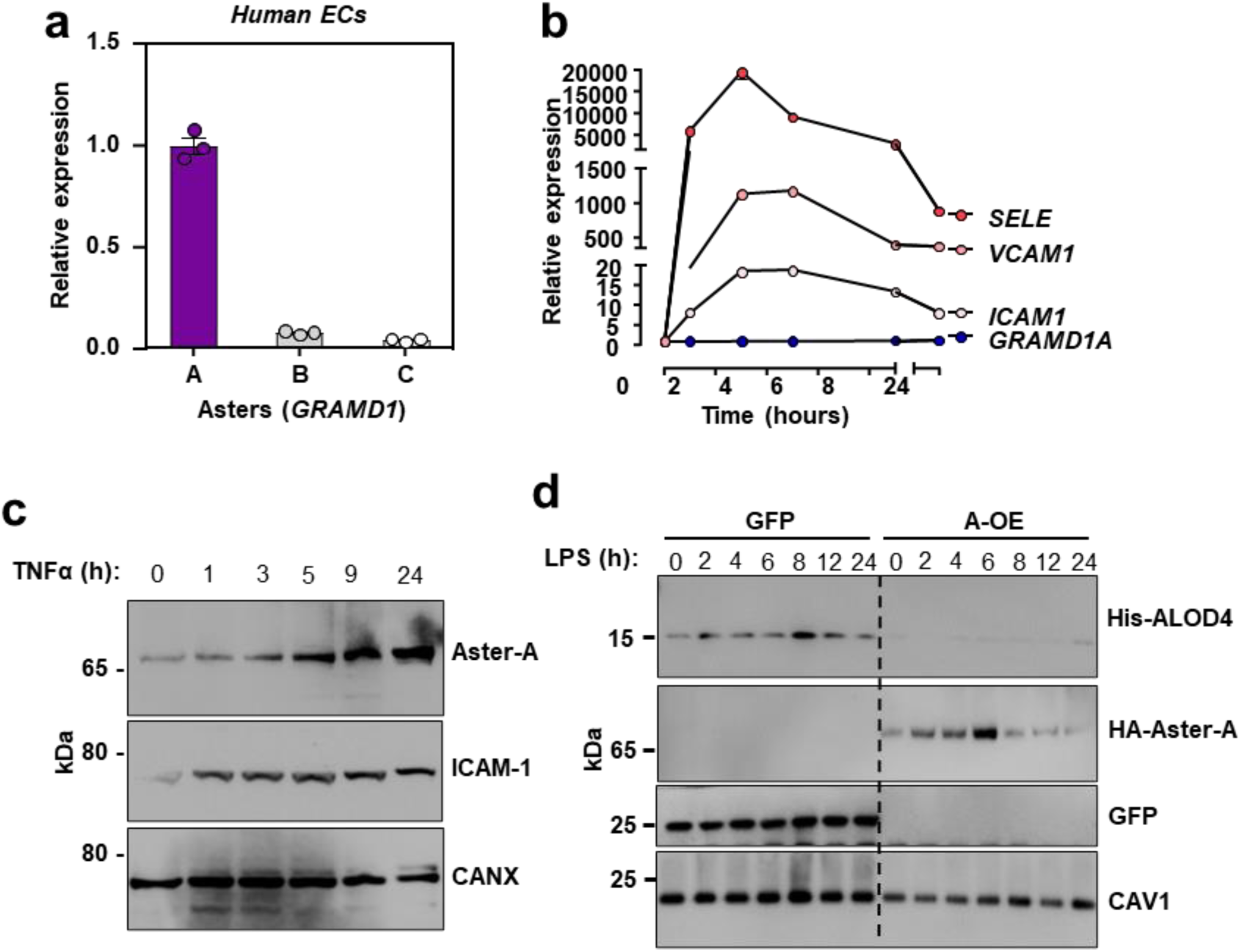
Aster-A undergoes post-translational stabilization in ECs after activation. (a) *GRAMD1A* (Aster-A), *GRAMD1B* (Aster-B) and *GRAMD1C* (Aster-C) expression relative to *36B4* in primary HAECs. (b) qPCR for *SELE*, *VCAM1*, *ICAM1* and *GRAMD1A* (Aster-A) in immortalized HAECs treated with TNFα for 0-24 h. (c) Endogenous Aster-A and ICAM-1 protein levels in HAECs treated with TNFα (10 ng/ml) for 0-24 h. (d) Western blots for His-ALOD4 and HA-Aster-A in HAECs stably overexpressing HA-Aster-A or GFP after LPS exposure for 0-24 h. Data are represented as mean ± SEM.

**Extended Data Fig. 5.**
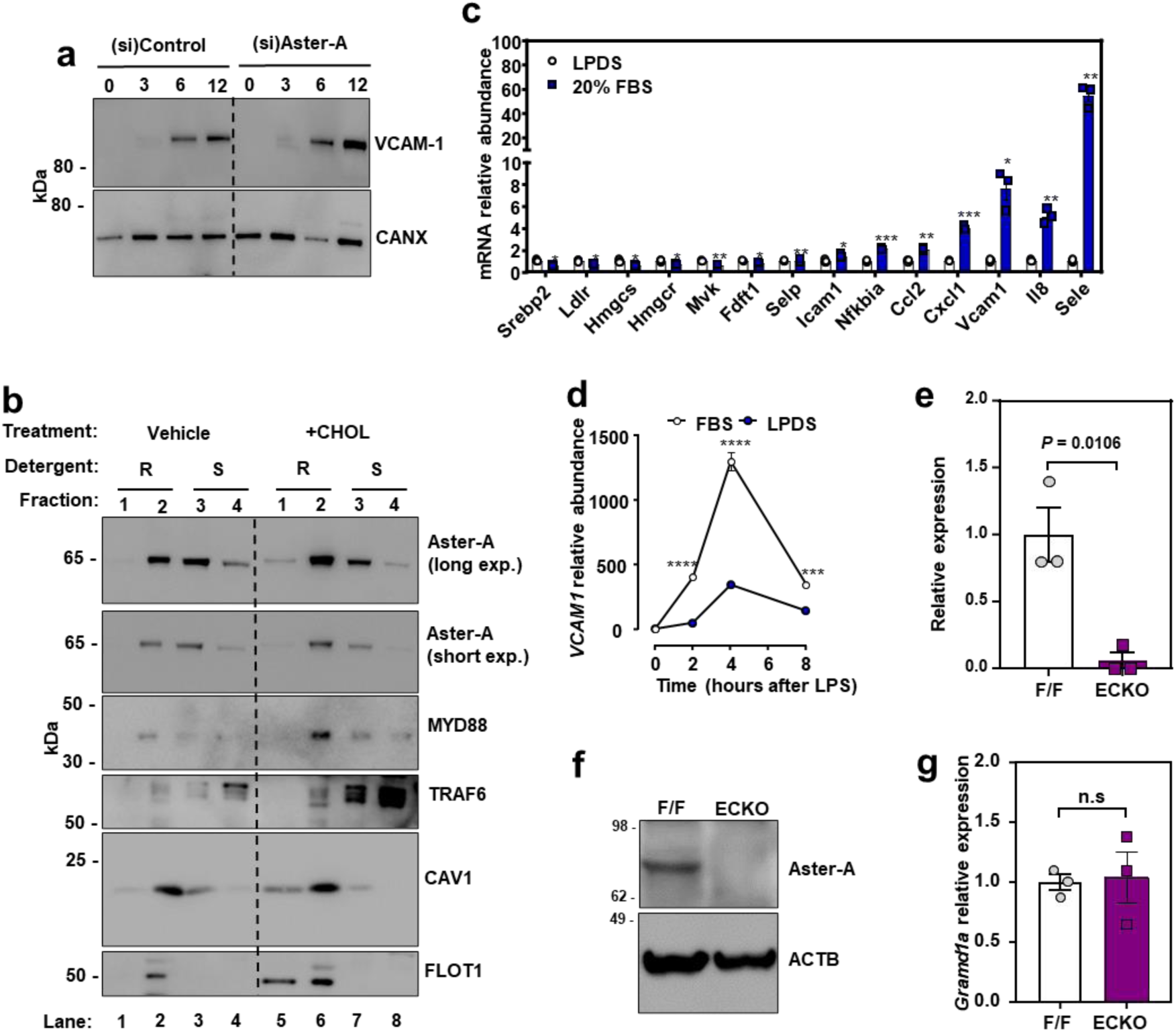
Nonvesicular cholesterol transport regulates inflammatory signaling in ECs. (a) Western blots for VCAM-1 in HAECs treated with (si)Control or (si)Aster-A and cultured in media containing 5% FBS with simvastatin overnight before exposure to LPS for the indicated times. (b) Western blots in HUVECs incubated with or without 100 µM MBCD-cholesterol for 1 h before fractionation of cells into Triton-X100 resistant or Triton-X100 soluble domains. (c) qPCR analysis in cells cultured in LPDS with simvastatin overnight before being loaded with or without the same media containing 20% FBS for 4 h. Target genes were normalized relative to *36B4*. (d) qPCR for *VCAM1* relative to *36B4* in HAECs cultured in either 10% FBS or 1% LPDS with simvastatin overnight before being stimulated with LPS (100 ng/ml) for 0-8 h. (e) *Gramd1a* mRNA in isolated hepatic ECs from F/F or ECKO mice. (f) Western blot for Aster-A in isolated hepatic ECs from F/F and ECKO mice. (g) *Gramd1a* mRNA in liver tissue from F/F or ECKO mice. n = 3 F/F and 3 ECKO. Data are represented as mean ± SEM.

**Extended Data Fig. 6.**
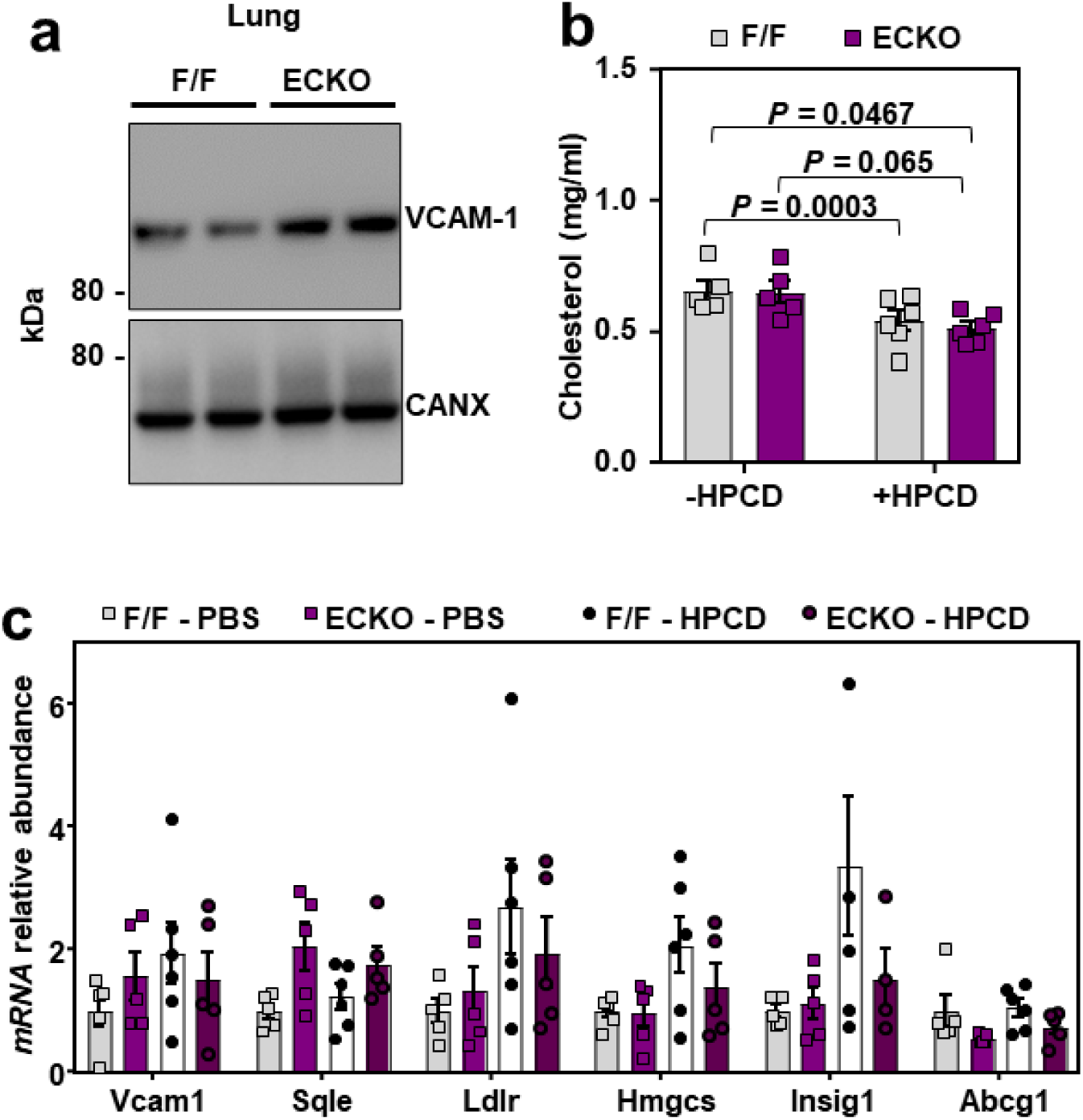
HPCD infusions lower VCAM-1 in response to LPS *in vivo*. (a) Western blots for VCAM-1 in the lungs of male F/F and ECKO mice 3 weeks after Cre induction. (b) Total plasma cholesterol in male F/F and ECKO mice injected with LPS (60 µg/mouse) for 20 mins before receiving i.v infusions of saline or HPCD (60 mg/mouse). Blood and tissues were collected 3 h after LPS injections. n = 5 F/F + saline, 5 ECKO + saline, 6 F/F + HPCD and 6 ECKO + saline. (c) qPCR in the hearts of male F/F and ECKO mice injected with LPS (60 µg/mouse) for 20 mins before receiving i.v infusions of saline or HPCD (60 mg/mouse). Tissues were collected 3 h after LPS injections. n = 5 F/F + saline, 5 ECKO + saline, 6 F/F + HPCD and 6 ECKO + saline. Data are represented as mean ± SEM with individual mice noted as dots.

## Acknowledgments

Confocal microscopy was performed at the California NanoSystems Institute of Advanced Light Microscopy/Spectroscopy Facility at UCLA. TIRF microscopy was performed at Biomedicum Imaging Unit supported by University of Helsinki/HiLIFE and Biocenter Finland. This work was supported by NIH grants DK126779, HL146358 and a Foundation Leducq Transatlantic Network of Excellence (19CVD04). J.P. K. was supported by an American Heart Association postdoctoral fellowship (903306). X.X. was supported by an American Heart Association postdoctoral fellowship (18POST34030388). Y.G. was supported by a Damon Runyon Cancer Research Foundation and Mark Foundation postdoctoral fellowship (DRG2424-21). S.H. was supported by a Jim Easton CDF Investigator award; A.F. was supported by a Ermenegildo Zegna Founder’s Scholarship (2017) and by an American Diabetes Association postdoctoral fellowship (1-19-PDF-043-RA). R.T.N. was supported by a T32GM008042 grant to the UCLA-Caltech Medical Scientist Training Program. A.N. was supported by a grant from the NIDDK (T32DK007180). J.J.M. was supported by American Heart Association Career Development Award 19CDA34760007. K.B was supported by NIH DP2 (GM146246-02) and the David and Lucile Packard Foundation Packard Fellowship. L.V. was supported by Instrumentarium Science Foundation and The Finnish Medical Foundation. E.I. was supported by Jane and Aatos Erkko Foundation and Sigrid Juselius Foundation.

## Author Contributions

J.P.K, X.X and P.T. contributed conceptualization; J.P.K., X.X, S.H., M.V., L.V., E.I., K.B., J.J.M. contributed methodology; J.P.K., X.X., Y.G., S.K., S.H., M.V., A.F., L.V., A.N., R.T.N., M.J.T contributed investigation; J.P.K., X.X. and P.T. contributed writing the manuscript; P.T. contributed funding acquisition; M.J.L., E.I., K.B., J.J.M., P.T. contributed resources; P.T. contributed supervision.

